# Construction and rescue of a DNA-launched DENV2 16681 infectious clone

**DOI:** 10.1101/2022.08.26.505435

**Authors:** Madeline Holliday, Lochlain Corliss, Nicholas J. Lennemann

**Affiliations:** Department of Microbiology, University of Alabama at Birmingham, Birmingham, AL, USA

**Author notes:** Corresponding author: Nicholas J. Lennemann, Department of Microbiology, University of Alabama at Birmingham, 845 19^th^ St S, BBRB 409B, Birmingham, AL 35294.

**Keywords:** positive-strand RNA virus, flavivirus, dengue virus, infectious clone, reverse genetics

## Abstract

Flaviviruses represent a large group of globally significant, insect-borne pathogens. For many of these viruses, there is a lack of antivirals and vaccines. Thus, there is a need to continue the development of tools to further advance our efforts to combat these pathogens, including reverse genetics techniques. Traditionally, reverse genetics methods for flaviviruses rely on producing infectious RNA from *in vitro* transcription reactions followed by electroporation or transfection into permissive cell lines. However, production of Zika virus has been successful from CMV-promoter driven expression plasmids, which provides cost and time advantages. In this report, we describe the design and construction of a DNA-launched infectious clone for dengue virus (DENV) serotype 2 strain 16681. An artificial intron was introduced in the nonstructural protein 1 segment of the viral genome to promote stability in bacteria. We found that rescued virus maintained similar replication kinetics in several cell lines commonly used in flavivirus research. Thus, we present a rapid and cost-effective method for producing DENV2 strain 16681 from plasmid DNA. This construct will be a useful platform for the continued development of anti-DENV therapeutics and vaccines.

## INTRODUCTION

Flaviviruses are a large group of insect-borne viruses that represent a significant risk to human health throughout most of the world. Dengue virus (DENV) is transmitted through the bite of infected mosquitos in tropical and sub-tropical regions [1,2]. More than 3 billion people live in regions that put them at risk of being infected with DENV, making it the most prevalent insect-borne virus in the world [3,4]. Disease from DENV infection can range from mild, febrile illness to hemorrhagic fever or shock that can lead to death [5]. To date, there are no approved antivirals for DENV, and the vaccine is not approved for use in children and elderly, which are populations most at risk[6,7]. Thus, there is a need to continue to develop tools that can be used to advance our efforts in antiviral and vaccine research.

Reverse genetics systems are important tools for virology research because they help to maintain viral genome sequence integrity and provide a platform to study specific mutations. Traditionally, flaviviruses have been rescued from cDNA clones expressed from a plasmid using *in vitro* transcription [8–15]. However, this can be a time-consuming process. Several flaviviruses have been successfully rescued from cytomegalovirus (CMV) promoter-driven plasmids after direct transfection of permissive cells [16–20]. A common issue observed during the production of flavivirus cDNA clones is the instability of the genome in bacteria due to the presence of cryptic promoters [17,21]. This was overcome for ZIKV cDNA clones by introducing synthetic introns into the genomic sequence, which force the viral polyprotein open reading frame (ORF) out of frame [19,20]. Alternatively, DENV has been successfully rescued from a bacterial artificial chromosome (BAC) system [22]. However, these systems can be complicated and time-consuming to produce. In this report, we describe the construction and rescue of DENV serotype 2 strain 16681 (DENV2) from a CMV promoter-driven cDNA clone. We found that introduction of a synthetic intron into NS1 stabilized the plasmid in bacteria and that virus was efficiently rescued from transfected cells using a cost-effective transfection reagent. Further, the rescued virus maintained many of the characteristics of the parental virus. Together, we show that this system provides a rapid and cost-effective alternative to the traditional RNA-launched cDNA infectious clones.

## MATERIALS AND METHODS

### Cell culture

Human embryonic kidney (HEK) 293T cells, A549 cells, U2OS cells, and C6/36 cells were maintained in Dulbecco’s modified Eagle’s media (DMEM) supplemented with 10% fetal bovine serum (FBS) and 100 U/mL penicillin/streptomycin (P/S). Human hepatoma cells (Huh7) were maintained in DMEM supplemented with 10% FBS, 9 g/L glucose, and 100U/mL P/S. VeroE6 cells (a gift from Dr. Kevin Harrod, University of Alabama at Birmingham) were maintained in Modified Eagle’s media (MEM) supplemented with 10% FBS and 100U/mL P/S. Baby hamster kidney (BHK-21) cells were maintained in MEM supplemented with 10% FBS, 1x non-essential amino acids, 1x sodium pyruvate, and 100U/mL P/S. All mammalian cells were maintained in humidified incubators at 37 °C. C6/36 cells were maintained in humidified incubators at 28 °C.

### Construction of infectious clone plasmid

Figure 1 and Table 1 summarize the plasmid design and construction of pcDNA6.2 DENV2 16681. The vector was obtained from PCR using Vector-F and Vector-R primers with the pcDNA6.2 ZIKV-MR766-HDVr plasmid [20]. This PCR introduced homologous sequences corresponding to the 5’UTR (22nt) and 3’UTR (18nt) of DENV2 into the linearized vector. The sequence encoding the DENV2 5’UTR was obtained as a gBlock (IDT) flanked by sequences homologous to the vector (22nt) and PCR A fragment (24nt). PCR A was obtained from DENV2 16681 CprME plasmid (a gift from Carolyn Coyne, Duke University) using PCR-A_F and PCR-A_R [23]. The synthetic intron was obtained as a gBlock (IDT) flanked by homologous sequences to PCR A (43nt) and the PCR B (20nt). PCR B was amplified from the subgenomic DENV2 replicon pcDNA3.1 DENVrepGFP/Zeo (a gift from Carolyn Coyne, Duke University) using PCR-B_F and PCR-B_R, which introduced sequence homologous to the vector at the 3’ end (15nt) [23]. PCR products and gBlocks were assembled using NEB HiFi Assembly, according to manufacturer’s protocol. Reactions were transformed in Stbl2 competent cells (Invitrogen), according to manufacturer’s protocol, and grown at 30 °C. Full plasmid sequencing was performed by Plasmidsaurus.

**Figure 1.**
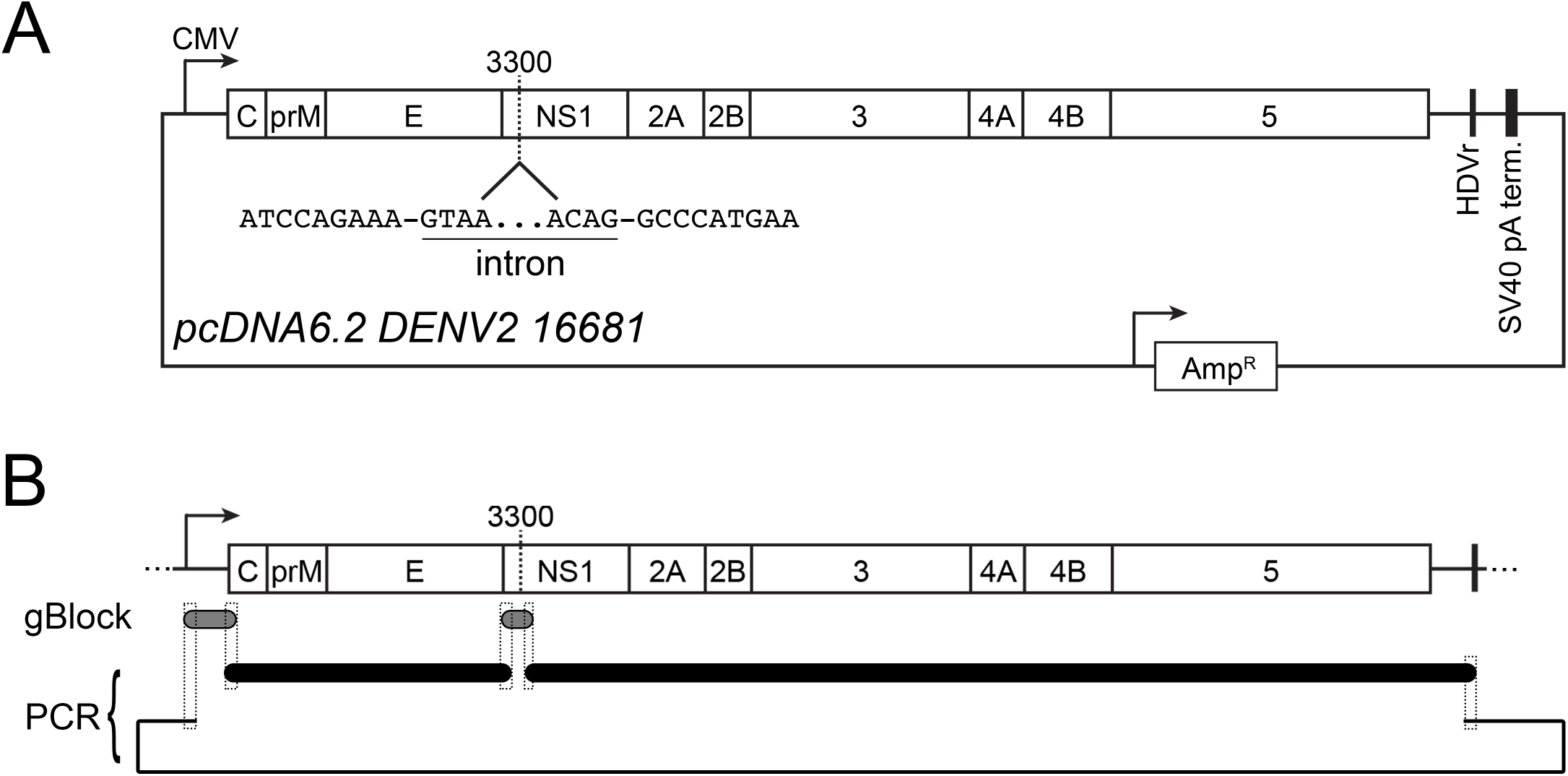
Design of pcDNA6.2 DENV2 16681. Schematics of a CMV promoter-driven cDNA clone of DENV2 strain 16681. (A) Plasmid map of pcDNA6.2 DENV2 16681. The viral genome lies between a CMV promoter and HDVr, followed by a SV40 polyA termination sequence. The location of the artificial intron is indicated at nucleotide 3300. Underlined sequence corresponds to the 5’ and 3’ ends of the intron. (B) Construction of pcDNA6.2 DENV2 16681. Overlapping regions between gBlocks (gray) and PCR products (black) are shown by dashed line boxes.

### Preparation of virus stocks

Laboratory stocks of DENV2 16681 (DENV2, a gift from Dr. Carolyn Coyne, Duke University) were prepared from C6/36 cells at 33 °C, as previously described. DENV2 was rescued from HEK 293T cells transfected with pcDNA6.2 DENV2 16681 plasmid using polyethylenimine (PEI, 25 kDa) at a 1:1 ratio of DNA (ug) to 1mg/mL PEI stock (µL) [24]. Virus containing supernatants were harvested at various times post transfection and assayed for infectious virus. Viral titers were determined by fluorescent focus forming unit assays, as previously described[25]. Briefly, 10-fold dilutions of virus containing supernatant were used to infect VeroE6 cells in 96-well plates. At ∼ 48hpi, cells were fixed in 4% paraformaldehyde diluted in PBS, permeabilized with 0.1% Triton 100-X diluted in PBS, and viral E protein was detected using cell culture derived monoclonal antibody 4G2 and fluorophore-conjugated secondary antibodies.

### Virus infections

Multicycle replication kinetics were determined in the indicated cell lines that were infected with 0.01 FFU/cell in a 6-well plate. At indicated times post infection, 100uL of supernatant was collected and stored at -80 °C until being titrated by FFU assay.

### Immunoblots

Lysates from transfected HEK 293T cells were lysed in 1x RIPA + protease inhibitor cocktail (Sigma). Lysates were clarified by centrifugation at 4 °C and 12,000 xg for 10 min. Clarified lysates were separated by SDS PAGE using a 4-20% Tris-glycine polyacrylamide pre-cast gel (BioRad) and transferred to nitrocellulose membranes. Following 30 min of blocking in PBS + 10% non-fat milk, membranes were probed using rabbit anti-DENV NS3 (1:2,000; GeneTex) and mouse anti-actin (1:10,000; ProteinTech) diluted in PBST + 5% BSA. Proteins were visualized using near infrared dye-conjugated secondary antibodies (LiCor) diluted in PBST + 5% non-fat milk and imaged on an Odyssey CLx imaging system (LiCor).

### Immunofluorescence microscopy

Transfected 293T cells seeded on poly-D-lysine coated 24-well plates were fixed at the indicated times post infection using PBS+ 4% paraformaldehyde for 10 min, permeabilized with PBS + 0.1% Triton 100-X for 10 min, washed in PBS, and incubated with mouse anti-dsRNA antibody (Kerafast) for 1h. Following primary antibody incubation, cells were washed in PBS x3 and incubated with AlexaFluor-conjugated secondary antibodies (Invitrogen). Following washing, samples were incubated with PBS + 300nM DAPI and imaged on a IX83 Olympus inverted fluorescent microscope using a 10x objective.

### Statistics

One-way ANOVA analyses were performed using Prism 9 Software (GraphPad).

## RESULTS AND DISCUSSION

### Design and construction of a DNA-launched DENV2 16681 infectious cDNA clone

To simplify production of DENV2, we sought to design a CMV-promoter driven infectious clone. We introduced the DENV2 16681 genome (NC_001474.2) into a modified pcDNA6.2 expression plasmid between the CMV promoter and a hepatitis delta ribozyme (HDVr), which self-cleaves upon transcription to maintain 3’ UTR sequence integrity (Figure 1A) [20]. Initial attempts to obtain clones containing the viral genome were unsuccessful due to large deletions starting in the nonstructural protein 1 (NS1) segment of the genome. Previous reports have demonstrated the instability of flavivirus genomes in plasmids due to the presence of cryptic promoters [17,21,26]. However, ZIKV has been successfully rescued via the insertion of a synthetic intron into the viral genome, which forces the viral ORF out of frame [19,20]. Thus, we took advantage of this strategy for the rescue of DENV2 clones by introducing an artificial intron at nucleotide 3300 of the DENV2 genome (Figure 1A). We successfully obtained a DNA-launched infectious clone through the assembly of gBlock DNA fragments, PCR-derived fragments of the viral genome, and a PCR-derived pcDNA6.2 vector containing the HDVr (Figure 1B and Table 1). Sequencing of recovered plasmid confirmed correct assembly of fragments containing no errors compared to the NC_001474.2 GenBank entry.

### Rescue of DNA-launched DENV2 16681

Next, we sought to rescue virus from our constructed plasmid. Supernatants from HEK 293T cells transfected with pcDNA6.2 DENV2 16681 plasmid showed the presence of infectious DENV2 starting at 48h post transfection (∼2 × 10^3^ FFU/mL). The amount of virus in the supernatant increased up to the final time point of 120h, when titers reached 2 × 10^6^ FFU/mL (Figure 2A). We further validated the production of DENV2 via immunoblot analysis of lysates from transfected HEK 293T cells. Consistent with our titer data, we observed an increase in nonstructural protein 3 (NS3) over time, with the peak expression occurring at 120h post transfection (Figure 2B). The presence of NS3 in cell lysates also indicates successful splicing of the recombinant genome, since expression of this protein is dependent on removal of the artificial intron. Furthermore, immunofluorescence shows an increase in dsRNA in transfected HEK 293T cells from 48 to 120h post transfection, which indicates the presence of DENV2 replication intermediates (Figure 2C). Together, these results show infectious DENV2 can be rescued from a CMV promoter-driven expression plasmid.

**Figure 2.**
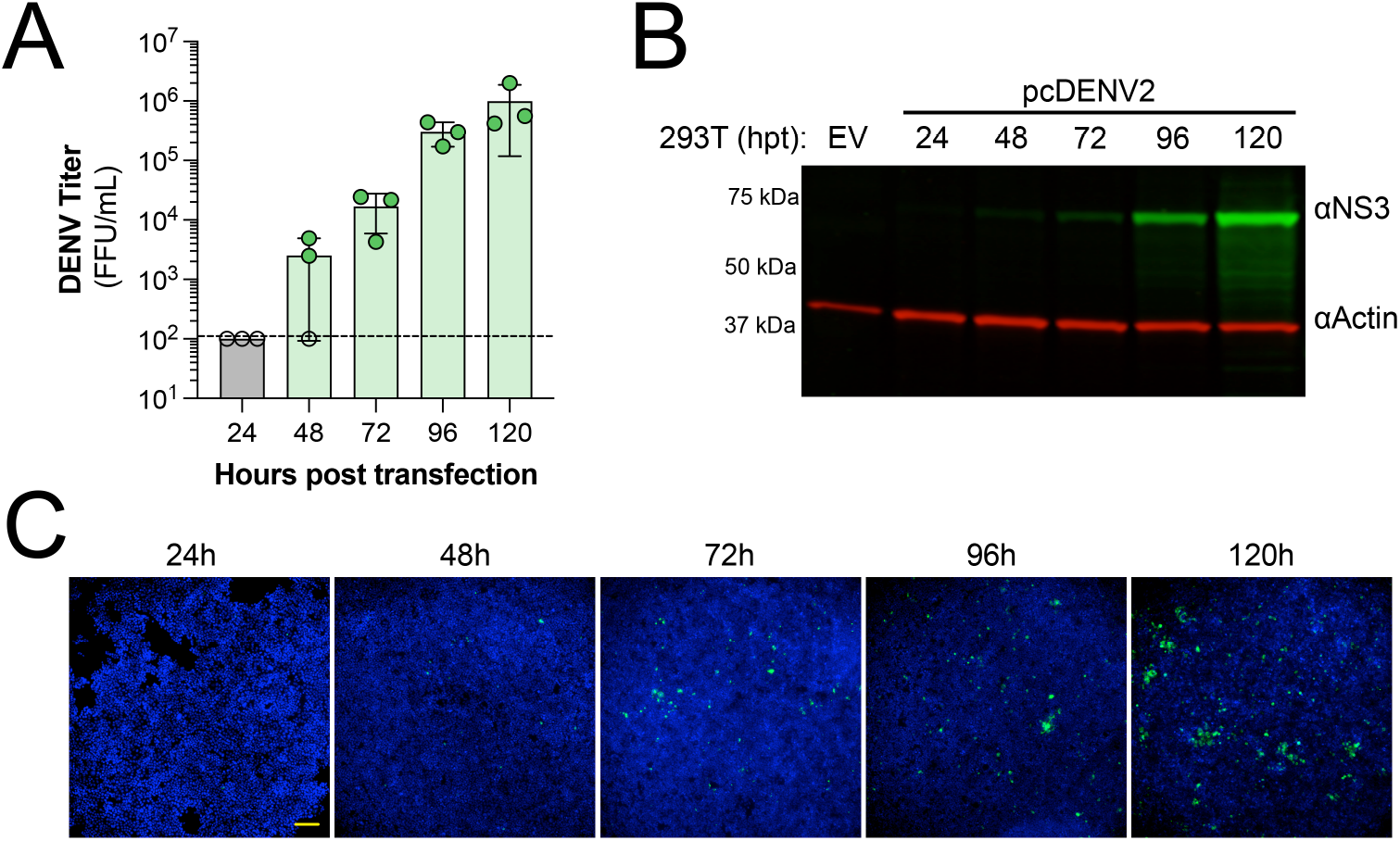
Rescue of DENV2. Validation of rescued virus. (A) Titer of DENV2 from pcDNA6.2 DENV 16681 transfected HEK 293T cells at indicated times post transfection. Data represents the average FFU/mL ± SD of three independent transfections. Open circles represent samples below the limit of detection, shown by the dashed line. (B) Representative immunoblot of lysates from transfected HEK 293T cells collected at the indicated hours post transfection (hpt). DENV NS3 is shown in green and actin is shown in red. Experiment was performed three independent times. (C) Immunofluorescence staining of transfected HEK 293T cells fixed at the indicated time post transfection. Viral dsRNA replication intermediate is shown in green and nuclei stained with DAPI are shown in blue. Scale bar represents X µm. Representative images are shown from one of three independent experiments.

### Characterization of rescued DENV2 16681

Next, we compared the replication kinetics of rescued DENV2 to parental DENV2 in various cell lines used in the literature. We performed multicycle replication assays in BHK-21, A549, VeroE6, U2OS, Huh7, and C6/36 cells from 0 to 72h post infection using rescued and stock virus. Overall, we observed similar replication kinetics in BHK-21, A549, VeroE6, and U2OS cells (Figure3A-D). At later time points, we observed higher titers in rescued virus from Huh7 cells compared to stock DENV2; however, this was not statistically significant (Figure 3E). Conversely, we observed significantly lower titer of rescued virus from C6/36 infected cells compared to stock virus at 72h post infection (Figure 3F). Differences in virus titer from infected Huh7 and C6/36 cultures could suggest the presence of tissue culture adapted mutants that are present in the stock parental virus compared to the rescued virus, which originated from a single genomic sequence. Overall, our results suggest rescuing DENV2 from pcDNA6.2 DENV 16681 is a rapid and cost-effective method for generating laboratory stocks of virus.

**Figure 3.**
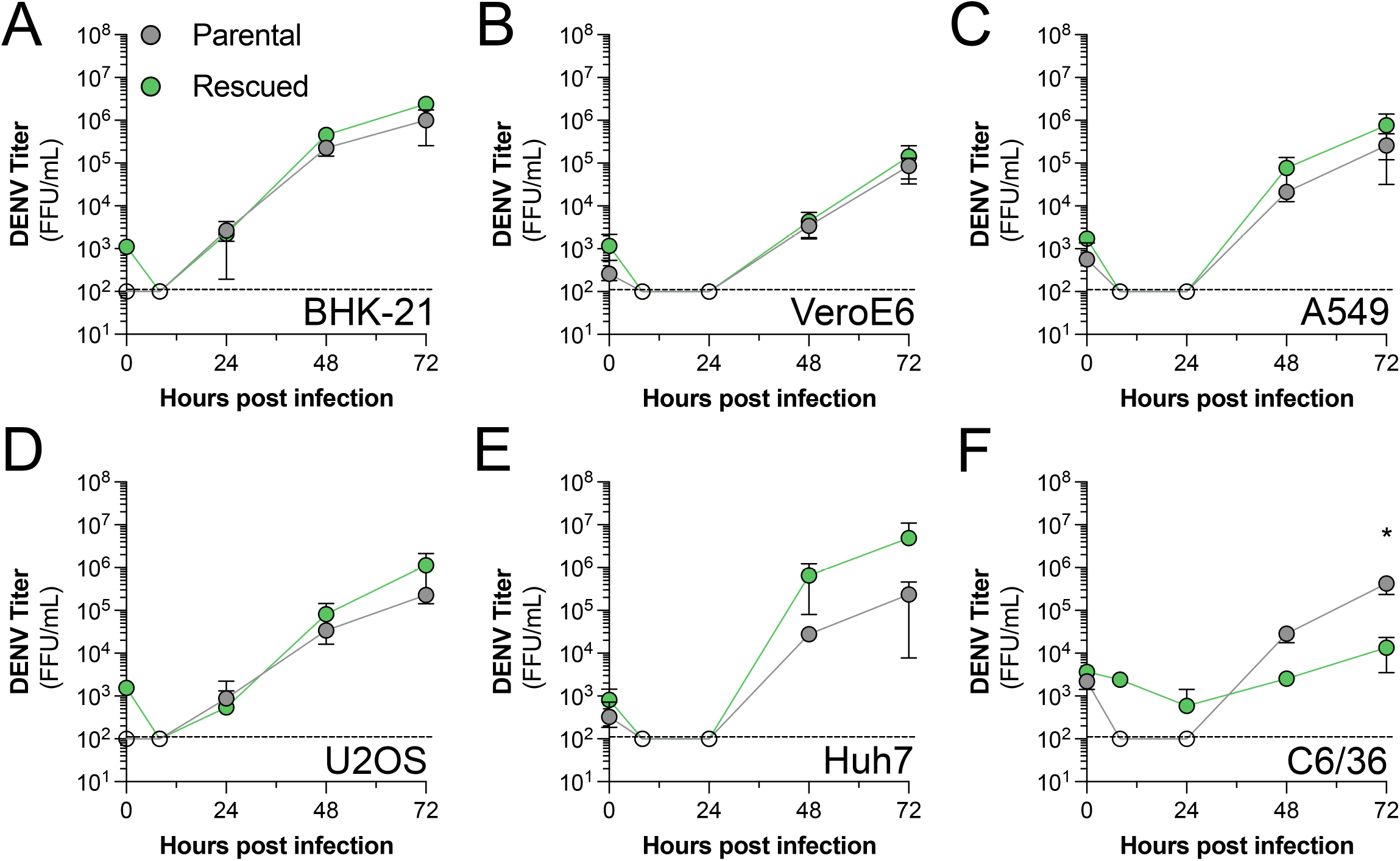
Replication kinetics of rescued virus in commonly used cell lines. (A-F) Indicated cell lines were infected with parental or rescued virus at an MOI of 0.01 for 72 h. Supernatants were collected from the indicated time points and titrated on VeroE6 cells. Data is shown as the average FFU/mL ± SD of three independent experiments. Significance was determined using a 2-way ANOVA with multiple comparisons, * p < 0.01.

## CONCLUSIONS

Historically, cDNA clones of flaviviruses have been rescued via in vitro transcription systems [21]. However, these systems can be time-consuming and expensive. Thus, we developed a CMV promoter-driven plasmid to rescue DENV2 strain 16681 via direct transfection, similar to plasmids reported for other flaviviruses [17,19,20]. Using this strategy, we were able to successfully rescue infectious DENV2 virus that resembles parental virus. This platform will streamline the production and analysis of DENV2 for a variety of applications, including the development of antiviral therapeutics and vaccines.

## Acknowledgements

We would like to thank Samaneh Mehri (University of Alabama at Birmingham) for critical reading of this manuscript. We would also like to thank Dr. Carolyn Coyne (Duke University), Dr. Guangxiang Luo (University of Alabama at Birmingham), and Dr. Kevin Harrod (University of Alabama at Birmingham), for generously providing reagents that made this work possible. The work described in this article was supported by the following funds: K22AI143963 (NL) and Development Funds from the University of Alabama at Birmingham (NL). The funders had no role in study design, data collection and analysis, decision to publish, or preparation of the manuscript.

